# Bridging Gaps in Soil Ecology: Metagenomic Insights into Microbial Diversity and Functionality Across Brazil’s Biomes

**DOI:** 10.1101/2025.05.16.654316

**Authors:** Luisa Mayumi Arake de Tacca, Rayane Nunes Lima, Patrícia Verdugo Pascoal, Marco Antônio de Oliveira, Carlos Alexandre Xavier de Azevedo, Deborah Bambil, Paula Mizotte, Grácia Maria Soares Rosinha, Daniela Matias de Carvalho Bittencourt, Diana Signor, Magna Soelma Beserra de Moura, José Pedro Pereira Trindade, Leandro Bochi da Silva Volk, Fernando Antônio Fernandes, Ricardo Lopes, Jean Luiz Simões-Araújo, Marcelo Freire, Elibio Rech

## Abstract

Microorganisms participate in complex interactions involving different kingdoms, so rhizosphere biodiversity mapping is essential for understanding how microbes interact with each other in the soil and with roots. Although soil microbial communities are remarkably diverse and technological advances have provided a high capacity to acquire reliable sequence data, unique microbial taxa in soil, root and rhizosphere samples remain poorly described. For the first time, we organized a consortium to collect soil samples covering all Brazilian biomes, providing a comprehensive and unprecedented view of soil microbial diversity. This understanding is critical, especially within the context of climate change, which affects plant physiology, root exudation and, consequently, the composition and functionality of soil microbial communities. The interactions between soil, roots and rhizosphere are influenced by evolutionary and adaptive forces and shape the production of microbial natural products, which exhibit great therapeutic potential and Mapping and studying rhizosphere microbial biodiversity not only increases our knowledge of soil ecology but also offers valuable insights for developing sustainable practices. We employed both 16S/18S/ITS amplicon and metagenomic short-read shotgun sequencing methods to examine and catalogue the large-scale genomes of culture-independent rhizosphere microbes and their interactions with roots in six terrestrial Brazilian biomes, namely, the Amazon, Atlantic Forest, Cerrado, Caatinga, Pampa and Pantanal. Our results revealed the ubiquity of *Proteobacteria*, which reflects their adaptability to contrasting environments. Biomes with greater moisture availability, such as the Amazon and Pantanal, exhibited greater diversity and abundance of fast-growing bacteria, such as *Proteobacteria*, and nutrient cyclers, such as *Thaumarchaeota*. Arid and semiarid biomes, such as the Caatinga, were dominated by microorganisms tolerant to drought and nutrient-limited environments, such as *Actinobacteria. Acidobacteria*, which thrive in acidic, nutrient-poor soils, were very abundant in forest biomes. The *Planctomycetes* phylum also occurred more frequently in areas with a relatively high soil organic matter content, such as the Cerrado. *Bacteroidetes* was significantly more abundant in Pampa than in the other biomes. The results provide comprehensive insights into soil, root and rhizosphere biodiversity and not only enhance the knowledge of the fundamental biological processes sustaining plant life but also constitute a reliable sequencing databank to address present-day agricultural and environmental challenges.

## Background

Microbiome metagenomics has emerged as an effective tool for elucidating the complex and diverse microbial communities that inhabit various ecosystems. In Brazil, a country renowned for its unparalleled biodiversity and distinct biomes, understanding microbiome dynamics is necessary for comprehending ecosystem health, biodiversity conservation, and potential applications in various fields. The Brazilian biomes, encompassing the Amazon, Atlantic Forest, Cerrado (savanna), Caatinga, Pampa and Pantanal (wetland), exhibit a mosaic of climate zones and ecological niches, each demonstrating unique microbial diversity [1, 2]. The Amazon is the largest tropical forest in the world, covering a significant portion of the northern region of Brazil [2]. It is known for its unrivalled biodiversity and its vital role in regulating the Earth’s climate. Soil in the Amazon ranges from poor to fertile, supporting lush and unique vegetation [3]. The Atlantic Forest, which once stretched along Brazil’s Atlantic coast, has been significantly reduced under deforestation [4]. Nevertheless, it is recognized for its diverse ecosystems, including dense forests, mangroves and coastal dunes [5]. The Cerrado is a vast tropical savanna biome that covers much of central Brazil. Although severely affected by deforestation, it is one of the savannas with the greatest biodiversity globally and encompasses pastures, shrublands and forests, with different soils, from fertile to the poorest [6]. The Caatinga, a dry forest biome in Northeast Brazil, with the lowest rainfall, contains drought-adapted vegetation, with thorny plants and seasonal rainfall patterns, defying arid conditions and the presence of often less fertile soils [7]. Pampa, which stretches across southern Brazil, is known for its vast grasslands and fertile soil, supporting diverse agriculture and an ecosystem unique to the region [8]. The Pantanal, the largest humid tropical area worldwide and located mainly in western Brazil, exhibits high aquatic and terrestrial biodiversity and abundant seasonally flooded soils [9].

Each biome exhibits unique soil properties, water availability, plant species, and microbial communities. Comparative analysis of soil, root and rhizosphere samples is critical for advancing our understanding of soil–plant–microorganism interactions and nutrient cycling and absorption [10-12]. The rhizosphere, which constitutes a dynamic interface between roots and the surrounding soil, is enriched with root exudates, creating a microhabitat that fosters various types of interaction, with a diversity symbiotic microbial community [13]. Fungi, particularly arbuscular mycorrhizal fungi (AMF), form symbiotic relationships with plant roots, increasing micronutrient absorption, especially phosphorus and nitrogen, through an expanded hyphal network [14]. Moreover, there exists an association between roots and prokaryotes that plays a critical role in the absorption of key nutrients, such as nitrogen, carbon and phosphorous [15]. In nutrient-poor soils, such as the Cerrado or Caatinga, such symbioses are essential for plant survival [16-18]. In flooded environments such as the Pantanal, anaerobic microorganisms are fundamental for methane production and consumption under anoxic conditions. In dry environments such as Caatinga, the rhizosphere is crucial for water retention and carbon cycling through the decomposition of organic matter by fungi and bacteria. These interactions are complex and directly influence the productivity and sustainability of terrestrial ecosystems. In this way, investigating these interactions in tandem with the surrounding soil matrix provides a more comprehensive understanding of how microbial communities are distributed to mediate biogeochemical cycles, influence soil structures and optimize nutrient availability.

Recent advancements in high-throughput sequencing technologies have facilitated comprehensive investigations into the structures and functions of microbial communities within these biomes [19, 20]. Metagenomic approaches allow researchers to analyze the collective genetic material of microbial populations directly from environmental samples, offering insights into the functional potential and interactions of these communities. This has created new avenues for studying the role of microorganisms in plant-microbe interactions and biogeochemical cycling processes in Brazilian ecosystems. Several studies have contributed to the growing body of knowledge on microbiome metagenomics in Brazilian biomes, revealing novel microbial taxa and functional genes and highlighting their role in sustaining plant health and ecosystem resilience [21-38]. Here, we conducted a large-scale profile study and analyses microbial community structure by sequencing 16S/18S/internal transcribed spacer (ITS) rRNA gene amplicons and conducting shotgun sequencing to elucidate the complex web of microbial life in soil, root and rhizosphere samples that underpins the ecological resilience and sustainability of these unique ecosystems.

Through a combination of cutting-edge research and technological advances, this exploration advances our understanding of the intrinsic relationships between microorganisms and their environments, thereby paving the way for the formulation of informed conservation strategies and innovative applications across disciplines. Finally, this integrated approach offers valuable insights into ecosystem functioning, sustainable land-use practices, agricultural management and environmental conservation.

## Material and methods

### Sample collection

A total of 79 samples were collected, with the following distributions across the various biomes: Amazon (12), Atlantic Forest (6), Caatinga (10), Cerrado Chapada dos Veadeiros (CV) (24), Cerrado Parque Nacional de Brasília (PNB) (4), Pampa (14) and Pantanal (9). The collected samples were divided into soil, root and rhizosphere components. The selection of sampling locations was strategically executed to encompass a broad spectrum of ecological sites, facilitating an in-depth exploration of potential variations in microbial communities across these regions. These biomes exhibit contrasting vegetation and microclimatic conditions, ranging from semi-arid environments to lush, humid rainforests.

The soil samples were manually sieved to remove rocks and roots to generate uniform samples. The rhizosphere was subsequently obtained by washing the roots with a 1X phosphate-buffered saline (PBS) solution (pH: 8.3). The roots were carefully removed, and the samples were centrifuged at 5000 rpm for 10 minutes.

### DNA extraction

The total genomic DNA of the soil and rhizosphere samples was extracted from 250 mg of each sample via the DNeasy PowerSoil Kit (QIAGEN, Germany) following the manufacturer’s instructions. The plant roots were rinsed in a sterile PBS solution (pH: 8.3) to remove the rhizosphere, disinfected with a solution of 1% sodium hypochlorite with 0.05% Tween 20 and agitated gently for 3 minutes to remove the remaining contaminants. The plant roots were then immersed in 70% ethanol for 2 minutes, followed by thorough rinsing with sterile water. Afterwards, the roots were ground with liquid nitrogen, followed by total genomic DNA extraction using the DNeasy Plant Kit (QIAGEN, Germany) following the manufacturer’s instructions. The quality and quantity of DNA were determined by measuring the absorbance at 260/280 nm (A260/A280) on a NanoDrop device (Thermo, Massachusetts, USA) and an Invitrogen Qubit 4 fluorometer device (Thermo, Massachusetts, USA), respectively. DNA integrity was verified by 0.8% agarose gel electrophoresis.

### 16S/18S/ITS rRNA gene amplicon library and sequence data analysis

For each biome, we pooled the samples into soil, root and rhizosphere samples, resulting in a total of 21 diverse samples. These samples were subsequently sequenced to obtain their 16S, 18S and ITS regions.

The following primers were used to amplify the 16S V4 region:

515F GTGCCAGCMGCCGCGGTAA and 806R GGACTACHVGGGTWTCTAAT

The following primers were used to amplify the 18S V4 region:

528F GCGGTAATTCCAGCTCCAA and 706R AATCCRAGAATTTCACCTCT

The following primers were used to amplify the ITS region:

ITS1-1F-F CTTGGTCATTTAGAGGAAGTAA and ITS1-1F-R GCTGCGTTCTTCATCGATGC

Amplification and amplicon sequencing, as well as pooled shotgun sequencing, were performed at Novogene.

### Shotgun sequencing and data processing

For all the biomes, the samples were normalized and pooled. All the root, rhizosphere and soil samples for any given biome were merged into a single biome sample. We analysed samples on a gel to determine degradation. Fifty nanograms of each biome pool were sequenced using Illumina shotgun sequencing at Novogene.

To obtain shotgun sequencing data, the samples were processed and cleaned at Novogene. Cleaned sequencing files were paired, and a paired library was created for each biome. These libraries were employed to classify the microbes using Kaiju through standard settings on the KBase platform [39].

To obtain amplicon-sequencing data, the DADA2 pipeline was adopted to process clean reads [40], leading to the formation of OTUs. Taxonomic classification of each representative read, and OTU was conducted using the ribosomal database project (RDP) classifier within the SILVA database for bacterial species (with a confidence level of 70%) and the UNITE database for fungal species [41, 42]. OTU analysis included the determination of the relative abundance at both the genus and phylum levels. The analysis was performed via the Phyloseq package [43].

## RESULTS

### Sequencing

To reveal the actual domain diversity of the soil, roots and rhizosphere, metagenomic DNA was isolated, sampled, and subsequently sequenced through Illumina amplicon and shotgun sequencing of the V3/V4 region of 16S rRNA, V4 region of 18S rRNA and ITS ITS1-1F. Thus, via the use of this comprehensive approach in which both amplicon sequencing and shotgun methodologies are combined, we ensured the precision of the estimated relative gene category abundances. This involved the random subsampling of reads to match the sample with the lowest read count, a prerequisite for the subsequent downstream analyses (Fig. 1A).

**Fig. 1.**
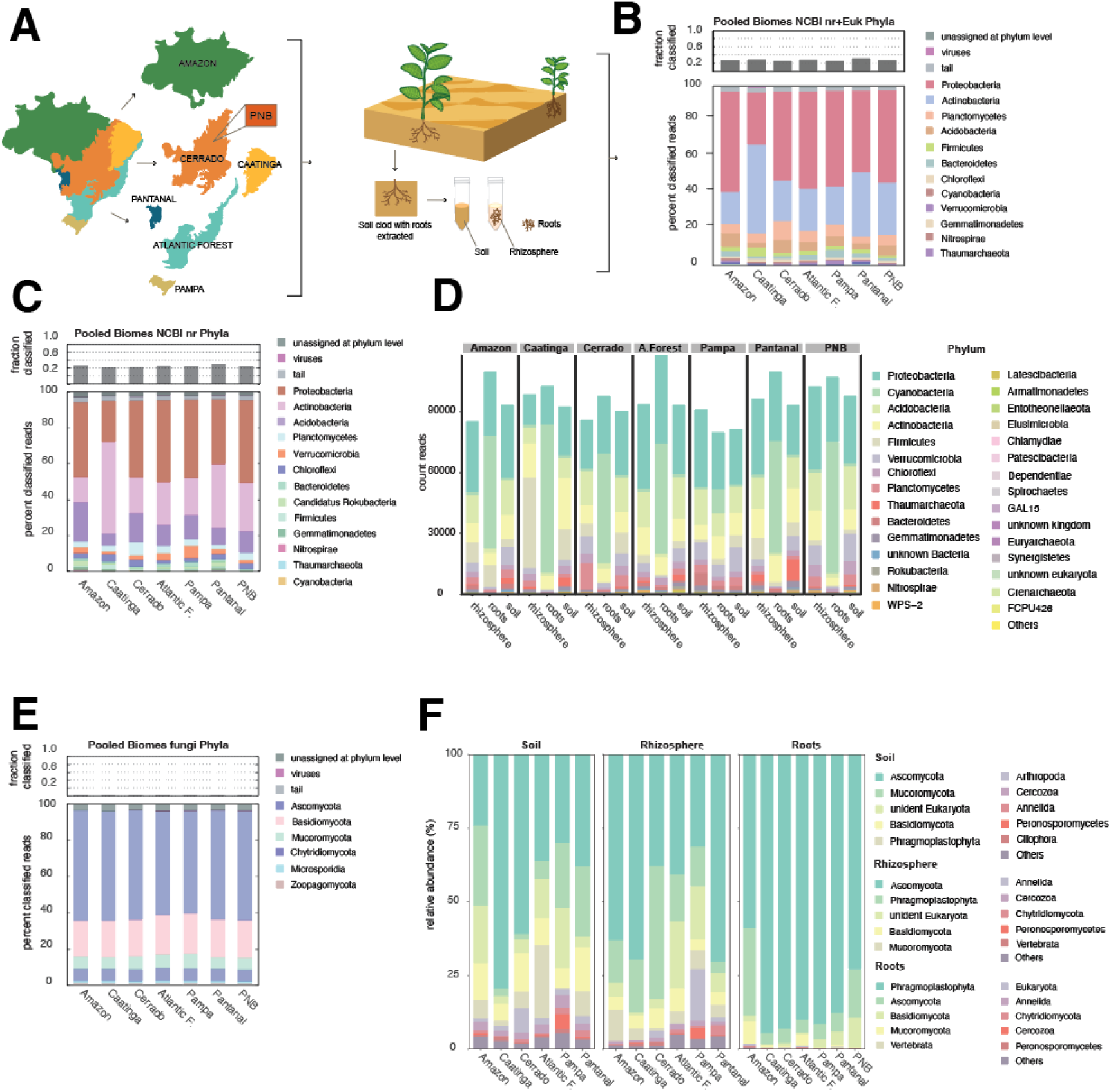
Sampling and sequencing of the Brazilian biomes. **A** Graphical representation of the Brazilian biomes sampled in this study. PNB stands for “Parque Nacional de Brasilia” and it is located within the Cerrado Biome. Each sample was collected for soil, roots and rhizosphere and sequenced through shotgun NGS and amplicon-seq using amplified 16S and 18S regions. **B** Shotgun data was classified using Kaiju for the phyla of each biome (NCBI nr+euk database). **C** Shotgun data was classified using Kaiju for the phyla of each biome (NCBI nr). **D** 16S regions of the biomes soil, roots and rhizosphere were amplified and sequenced with Illumina. Classification was done using the silva_nr_v132_train_set. **E** Shotgun data was classified using Kaiju for the phyla of fungi. **F** 18S regions of the biomes soil, roots and rhizosphere were amplified and sequenced with Illumina. Classification was done using the silva_nr_v138_train_set. Figures B, C and E were made at KBase using the Kaiju package and figures D and F were made in R using ggbartax in the MicrobiotaProcess R package.

To capture the overall biome-level representation, all individual samples from each biome were pooled into a single-biome source sample. These biome source samples were subjected to shotgun sequencing, resulting in a cumulative total of 72,569,233 base pairs (Table 1). This comprehensive sampling and sequencing strategy aimed to provide a robust foundation for elucidating the complex microbial dynamics in distinct Brazilian biomes.

**Table 1.** Summary of the sequencing data of the biomes of Brazil.

| **Biome** | **Number of paired-end reads** | **Bacterial reads (\*N, %)** | **Archaea reads (N,%)** | **Viral reads (N, %)** | **Fungi reads (N, %)** |
| --- | --- | --- | --- | --- | --- |
| **Amazon** | 49,824,684 | 1,137,692 | 9,809 | 8,532 | 53,572 |
| **Atlantic Forest** | 41,664,538 | 923,693 | 5,327 | 7,630 | 41,156 |
| **Caatinga** | 45,631,994 | 923,439 | 5,642 | 7,603 | 48,584 |
| **Cerrado_CV** | 35,895,044 | 738,355 | 2,699 | 5,579 | 38,008 |
| **Cerrado_PNB** | 62,818,310 | 1,356,564 | 2,557 | 9,974 | 64,363 |
| **Pampa** | 62,191,666 | 1,359,808 | 2,041 | 10,332 | 63,578 |
| **Pantanal** | 45,605,150 | 1,117,556 | 8,540 | 8,498 | 54,838 |

\*From Krona plots calculated with Kaiju

### Prokaryotic and eukaryotic read distributions across Brazilian biomes

The samples were normalized and pooled for each biome, and subsequently used for shotgun sequencing. Here, we could obtain a macroscopic overview of the microbes present in each Brazilian biome, while the amplicon-sequencing data provided a comprehensive view of prokaryotic and eukaryotic diversity across Brazilian biomes, given the separation between roots, soil and the rhizosphere. The differences in microbial community composition can be attributed to the unique ecological conditions in each biome, such as moisture, temperature, soil pH, soil nutrition and vegetation type. Below are detailed statistical annotated read results and phylum prevalence differences.

### Amazon forest

According to the analysis of the pooled shotgun data through Kaiju, at the phylum level, 698,227 and 5,249 reads were assigned to the *Bacteria* and *Archaea* domains, respectively. The most abundant bacterial sequences were *Proteobacteria*, accounting for 56% of the total sequences, followed by *Actinobacteria* (17.8%), *Acidobacteria* (7.4%), *Planctomycetes* (5.3%), *Bacteroidetes* (2.8%), *Firmicutes* (2.5%), *Chloroflexi* (1.2%), *Cyanobacteria* (1.1%) and *Nitrospirae* (1%). Moreover, the abundance levels of four phyla remained below 1%, namely, *Verrucomicrobia* (0.94%), *Gemmatimonadetes* (0.81%), *Deinococcus*-*Thermus* (0.28%), and *Spirochaetes* (0.14%). The abundance levels of 21 phyla remained below 0.1%, namely, *Chlorobi* (0.09%), *Chlamydiae* (0.08%), *Aquificae* (0.07%), *Thermotogae* (0.07%), *Armatimonadetes* (0.07%), *Thermodesulfobacteria* (0.06%), *Ignavibacteriae* (0.06%), *Rhodothermaeota* (0.05%), *Deferribacteres* (0.04%), *Kiritimatiellaeota* (0.04%), *Calditrichaeota* (0.03%), *Synergistetes* (0.03%), *Fusobacteria* (0.02%), *Tenericutes* (0.02%), *Candidatus Omnitrophica* (0.01%), *Elusimicrobia* (0.01%), *Dictyoglomi* (0.01%), *Atribacterota* (0.01%), *Balneolaeota* (0.01%), *Candidatus Bipolaricaulota* (0.01%), and *Chrysiogenetes* (0.01%). The least abundant phyla were *Candidatus Cloacimonetes* (0.006%), *Candidatus Saccharibacteria* (0.006%), *Fibrobacteres* (0.005%), *Caldiserica* (0.004%), *Coprothermobacterota* (0.003%) and *Candidatus Absconditabacteria* (0.0008%). The most abundant archaeal sequences were *Thaumarchaeota* (0.34%) and *Euryarchaeota* (0.31%), whereas the abundance levels of other phyla did not exceed 0.1%, including *Crenarchaeota* (0.05%), *Candidatus Thermoplasmatota* (0.02%), *Candidatus Korarchaeota* (0.003%), *Candidatus Micrarchaeota* (0.003%), *Candidatus Lokiarchaeota* (0.001%), and *Candidatus Nanohaloarchaeota* (0.0004%). A total of 201 reads (0.03%) were assigned to viral genomes, whereas unassigned and unclassified reads (1,787,557) accounted for 71% of the total reads on average (Fig. 1B and 1C, respectively). Notably, the pooled shotgun composition classified on the basis of the National Center for Biotechnology Information (NCBI) nr+euk database mostly matched that resulting from the amplicon-sequencing data classified on the basis of Silva’s training set (Fig. 1D). Like in the shotgun data, in the amplicon-sequencing data, *Proteobacteria* also dominated in the rhizosphere and soil. However, we observed a different pattern for the roots from the Amazon samples. Compared with those in the soil and rhizosphere, the roots were dominated by *Cyanobacteria*, followed by *Proteobacteria* and *Actinobacteria*. Notably, the relative abundance of *Firmicutes* was also greater than that in the soil and rhizosphere (Fig. 1E and 1D, respectively).

In the samples from the Amazon, 51,693 reads were assigned to fungi at the phylum level. The most abundant fungal sequence was *Ascomycota* (61%), followed by *Basidiomycota* (20%), *Mucoromycota* (7.7%), *Chytridiomycota* (6.5%), *Zoopagomycota* (1.3%) and *Microsporidia* (1.8%). At the genus level, the most prevalent taxa were *Aspergillus* (5.3%), *Trichoderma* (4%), *Fusarium* (2.5%), *Spizellomyces* (2.4%), *Lobosporangium* (2.1%), *Batrachochytrium* (2%), *Synchytrium* (1.7%), *Rhizophagus* (1.4%), *Penicillium* (1.4%) and *Phycomyces* (1.4%). These genera represent the top 10 dominant fungal populations identified. Unassigned and unclassified reads (2439541) accounted for 98% of the total reads on average.

### Caatinga

At the phylum level, 602,062 and 5,550 reads were assigned to the *Bacteria* and *Archaea* domains, respectively. The most abundant bacterial sequences were *Actinobacteria*, accounting for 49.5% of the total sequences, followed by *Proteobacteria*, at 29%; *Planctomycetes*, at 5.1%; *Firmicutes*, at 5.1%; *Acidobacteria*, at 2.3%; *Chloroflexi*, at 1.5%; *Bacteroidetes*, at 1.4%; and *Cyanobacteria*, at 1.1%. Moreover, the abundance levels of five phyla remained below 1%, namely, *Gemmatimonadetes* (0.97%), *Verrucomicrobia* (0.44%), *Deinococcus*-*Thermus* (0.38%), *Nitrospirae* (0.3%) and *Spirochaetes* (0.12%). Sixteen phyla exhibited abundance levels below 0.1%, namely, *Armatimonadetes* (0.06%), *Chlorobi* (0.06%), *Thermotogae* (0.05%), *Aquificae* (0.05%), *Thermodesulfobacteria* (0.05%), *Rhodothermaeota* (0.04%), *Chlamydiae* (0.04%), *Ignavibacteriae* (0.03%), *Deferribacteres* (0.03%), *Synergistetes* (0.03%), *Tenericutes* (0.02%), *Calditrichaeota* (0.02%), *Fusobacteria* (0.02%), *Kiritimatiellaeota* (0.02%), *Dictyoglomi* (0.01%) and *Candidatus Bipolaricaulota* (0.01%). The least abundant phyla were *Candidatus Saccharibacteria* (0.009%), *Balneolaeota* (0.009%), *Elusimicrobia* (0.008%), *Atribacterota* (0.008%), *Candidatus Omnitrophica* (0.008%), *Coprothermobacterota* (0.006%), and *Chrysiogenetes* (0.006%). *Candidatus Cloacimonetes* (0.004%), *Caldiserica* (0.004%), *Fibrobacteres* (0.003%), and *Candidatus Absconditabacteria* (0.0005%). The most abundant archaeal sequences were *Thaumarchaeota* (0. 49%) and *Euryarchaeota* (0.34%), whereas the abundance levels of the other phyla did not exceed 0.1%, including *Crenarchaeota* (0.04%), *Candidatus Thermoplasmatota* (0.01%), *Candidatus Lokiarchaeota* (0.001%), *Candidatus Korarchaeota* (0.001%), *Candidatus Micrarchaeota* (0.0006%), and *Candidatus Nanohaloarchaeota* (0.0001%). A total of 161 reads (0.02%) were assigned to viral genomes, whereas unassigned and unclassified reads (1673826) accounted for 73% of the total reads on average (Fig. 1B). In the amplicon-sequencing 16S data, the profile for the Caatinga biome slightly differed from the shotgun profile. In the latter, the dominant phylum was *Actinobacteria* (Fig. 1B and 1C). However, in the amplicon-sequencing 16S data, *Proteobacteria* was more dominant than *Actinobacteria* in the soil and rhizosphere.

For fungi, 46,630 reads were assigned at the phylum level. The most abundant fungal sequence was *Ascomycota* (60%), followed by *Basidiomycota* (20%), *Chytridiomycota* (6.5%), *Mucoromycota* (6.4%), *Microsporidia* (1.3%) and *Zoopagomycota* (1%). At the genus level, *Aspergillus* (6.3%), *Spizellomyces* (2.3%), *Fusarium* (2.1%), *Synchytrium* (2%), *Batrachochytrium* (2%), *Rhizophagus* (1.8%), *Lobosporangium* (1.6%), *Trichoderma* (1.4%), *Phycomyces* (1.4%), and *Talaromyces* (1.4%) were identified. These genera represent the top 10 dominant fungal populations identified. Unassigned and unclassified reads (2234969) accounted for 98% of the total reads on average (Fig.1E).

### Cerrado–Chapada dos Veadeiros (CV)

At the phylum level, 490,262 and 2,030 reads were assigned to the *Bacteria* and *Archaea* domains, respectively. The most abundant bacterial sequences were *Proteobacteria*, accounting for 50% of the total sequences, followed by *Actinobacteria* (22%), *Planctomycetes* (10%), *Acidobacteria* (7.1%), *Firmicutes* (2%), *Chloroflexi* (1.5%), *Bacteroidetes* (1.5%) and *Cyanobacteria* (1.1%). %). Moreover, the abundance levels of five phyla remained below 1%, namely, *Verrucomicrobia* (0.9%), *Gemmatimonadetes* (0.36%), *Deinococcus*-*Thermus* (0.27%), *Nitrospirae* (0.25%) and *Spirochaetes* (0.11%). The abundance levels of 14 phyla remained below 0.1%, namely, *Armatimonadetes* (0.07%), *Chlorobi* (0.07%), *Chlamydiae* (0.06%), *Ignavibacteriae* (0.05%), *Thermotogae* (0.05%), *Aquificae* (0.05%), *Thermodesulfobacteria* (0.04%), *Rhodothermaeota* (0.03%), *Kiritimatiellaeota* (0.03%), *Deferribacteres* (0.03%), *Synergistetes* (0.03%), *Calditrichaeota* (0.02%), *Fusobacteria* (0.02%) and *Tenericutes* (0.02%). The least abundant phyla were *Elusimicrobia* (0.009%), *Atribacterota* (0.009%), *Chrysiogenetes* (0.009%), *Candidatus Bipolaricaulota* (0.008%), *Dictyoglomi* (0.008%), *Balneolaeota* (0.007%), *Candidatus Omnitrophica* (0.007%), *Caldiserica* (0.005%), *Candidatus Saccharibacteria* (0.005%), *Fibrobacteres* (0.003%), *Coprothermobacterota* (0.003%), *Candidatus Absconditabacteria* (0.0014%) and *Candidatus Cloacimonetes* (0.0014%). The most abundant archaeal sequence was *Euryarchaeota* (0.27%), while the abundance levels of the other phyla did not exceed 0.1%, including *Thaumarchaeota* (0.07%), *Crenarchaeota* (0.03%), *Candidatus Thermoplasmatota* (0.02%), *Candidatus Micrarchaeota* (0.001%), *Candidatus Korarchaeota* (0.001%), *Candidatus Nanohaloarchaeota* (0.0006%) or *Candidatus Lokiarchaeota* (0.0006%). Eighty-four reads (0.03%) were assigned to viral genomes, whereas unassigned and unclassified reads (1302376) accounted for 72% of the total reads on average (Fig. 1B and 1C).

For fungi, 36,719 reads were assigned at the phylum level. The most abundant fungal sequence was *Ascomycota* (60%), followed by *Basidiomycota* (20%), *Mucoromycota* (7.2%), *Chytridiomycota* (6.6%), *Zoopagomycota* (1.1%) and *Microsporidia* (1.1%). At the genus level, *Aspergillus* (5%), *Spizellomyces* (2.5%), *Rhizophagus* (2.2%), *Batrachochytrium* (1.9%), *Lobosporangium* (1.8%), *Bacidia* (1.5%), *Exophiala* (1.5%), *Phycomyces* (1.5%), *Fusarium* (1.3%), and *Talaromyces* (1.2%) were identified. These genera represent the top 10 dominant fungal populations identified. Unassigned and unclassified reads (1758033) accounted for 98% of the total reads on average (Fig. 1E and 1F).

### Cerrado–Parque Nacional de Brasília (PNB)

At the phylum level, 907,146 and 2,831 reads were assigned to the *Bacteria* and *Archaea* domains, respectively. The most abundant bacterial sequences were *Proteobacteria* (accounting for 51% of the total sequences), *Actinobacteria* (28%), *Planctomycetes* (5.9%), *Acidobacteria* (5.7%), *Firmicutes* (1.5%) and *Bacteroidetes* (1.2%). The abundance levels of 6 phyla remained below 1%, namely, *Chloroflexi* (0.99%), *Cyanobacteria* (0.87%), *Verrucomicrobia* (0.74%), *Gemmatimonadetes* (0.33%), *Deinococcus*-*Thermus* (0.24%) and *Nitrospirae* (0.15%). Moreover, the abundance levels of 15 phyla remained below 0.1%, namely, *Spirochaetes* (0.08%), *Armatimonadetes* (0.05%), *Chlorobi* (0.05%), *Chlamydiae* (0.04%), *Thermotogae* (0.04%), *Aquificae* (0.03%), *Rhodothermaeota* (0.03%), *Thermodesulfobacteria* (0.03%), *Ignavibacteriae* (0.02%), *Kiritimatiellaeota* (0.02%), *Deferribacteres* (0.02%), *Calditrichaeota* (0.02%), *Synergistetes* (0.02%), *Fusobacteria* (0.01%) and *Tenericutes* (0.01%). The least abundant phyla were *Atribacterota* (0.008%), *Balneolaeota* (0.008%), *Dictyoglomi* (0.007%), *Elusimicrobia* (0.007%), *Candidatus Bipolaricaulota* (0.006%), *Chrysiogenetes* (0.005%), *Candidatus Omnitrophica* (0.005%), *Candidatus Saccharibacteria* (0.005%), *Caldiserica* (0.003%), *Fibrobacteres* (0.003%), *Candidatus Cloacimonetes* (0.002%) *Coprothermobacterota* (0.002%) and *Candidatus Absconditabacteria* (0.0004%). The most abundant archaeal sequence was *Euryarchaeota* (0.21%), while the abundance levels of the other phyla did not exceed 0.1%, including *Thaumarchaeota* (0.05%), *Crenarchaeota* (0.03%), *Candidatus Thermoplasmatota* (0.007%), *Candidatus Lokiarchaeota* (0.001%), *Candidatus Korarchaeota* (0.001%), *Candidatus Micrarchaeota* (0.0008%), and *Candidatus Nanohaloarchaeota* (0.0003%). A total of 157 reads (0.03%) were assigned to viral genomes, whereas unassigned and unclassified reads (2230781) accounted for 70% of the total reads on average (Fig. 1B and 1C).

For fungi, 61,996 reads were assigned at the phylum level. The most abundant fungal sequences were *Ascomycota* (60%), *Basidiomycota* (20%), *Mucoromycota* (6.6%), *Chytridiomycota* (6.4%), *Microsporidia* (1.1%) and *Zoopagomycota* (1%). At the genus level, *Aspergillus* (5%), *Spizellomyces* (2.3%), *Synchytrium* (2%), *Rhizophagus* (2%), *Batrachochytrium* (1.8%), *Lobosporangium* (1.7%), *Exophiala* (1.5%), *Fusarium* (1.4%), *Phycomyces* (1.3%) and *Bacidia* (1.3%) were identified. These genera represent the top 10 dominant fungal populations identified. Unassigned and unclassified reads (3078919) accounted for 98% of the total reads on average (Fig. 1E).

### Atlantic forest

At the phylum level, 619,723 and 3971 reads were assigned to the *Bacteria* and *Archaea* domains, respectively. The most abundant bacterial sequence was *Proteobacteria*, accounting for 54.2% of the total sequences, followed by *Actinobacteria* (26.3%), *Planctomycetes* (6%), *Acidobacteria* (5%), *Bacteroidetes* (2.7%) and *Firmicutes* (2.1%). *Moreover*, the abundance levels of seven phyla remained below 1%, namely, *Verrucomicrobia* (0.96%), *Cyanobacteria* (0.83%), *Gemmatimonadetes* (0.75%), *Chloroflexi* (0.71%), *Nitrospirae* (0.40%), *Deinococcus*-*Thermus* (0.24%), and *Spirochaetes* (0.10%). The abundance levels of 15 phyla remained below 0.1%, namely, *Chlorobi* (0.07%), *Armatimonadetes* (0.06%), *Chlamydiae* (0.05%), *Thermotogae* (0.05%), *Aquificae* (0.04%), *Rhodothermaeota* (0.04%), *Thermodesulfobacteria* (0.04%), *Ignavibacteriae* (0.04%), *Deferribacteres* (0.03%), *Kiritimatiellaeota* (0.03%), *Synergistetes* (0.03%) *Fusobacteria* (0.03%), *Calditrichaeota* (0.02%), *Tenericutes* (0.01%), and *Elusimicrobia* (0.01%). The least abundant phyla were *Balneolaeota* (0.009%), *Atribacterota* (0.008%), *Dictyoglomi* (0.007%), *Candidatus Omnitrophica* (0.007%), *Chrysiogenetes* (0.007%), *Candidatus Saccharibacteria* (0.006%), *Candidatus Bipolaricaulota* (0.006%), *Fibrobacteres* (0.005%), *Candidatus Cloacimonetes* (0.004%), *Coprothermobacterota* (0.003%), *Caldiserica* (0.003%), and *Candidatus Absconditabacteria* (0.0006%). The most abundant archaeal sequences were *Thaumarchaeota* (0. 36%) and *Euryarchaeota* (0.23%), whereas the abundance levels of the other phyla did not exceed 0.1%, including *Crenarchaeota* (0.03%), *Candidatus Thermoplasmatota* (0.007%), *Candidatus Korarchaeota* (0.001%), *Candidatus Micrarchaeota* (0.001%) and *Candidatus Lokiarchaeota* (0.0001%). A total of 330 reads (0.05%) were assigned to viral genomes, whereas unassigned and unclassified reads (1459532) accounted for 70% of the total reads on average (Fig. 1B and 1C).

For fungi, 39,525 reads were assigned at the phylum level. The most abundant fungal sequence was *Ascomycota* (57%), followed by *Basidiomycota* (22%), *Mucoromycota* (7.3%), *Chytridiomycota* (7.1%), *Microsporidia* (1.3%) and *Zoopagomycota* (1.1%). At the genus level, *Aspergillus* (4.8%), *Spizellomyces* (2.6%), *Trichoderma* (4%), *Batrachochytrium* (2.2%), *Lobosporangium* (2%), *Synchytrium* (2%), *Rhizophagus* (1.7%), *Phycomyces* (1.6%), *Fusarium* (1.3%) and *Rhizopus* (1.2%) were identified. These genera represent the top 10 dominant fungal populations identified. Unassigned and unclassified reads (2043701) accounted for 98% of the total reads on average (Fig. 1E).

### Pampa

At the phylum level, 853,361 and 2,933 reads were assigned to the *Bacteria* and *Archaea* domains, respectively. The most abundant bacterial sequence was *Proteobacteria*, accounting for 53% of the total sequences, followed by *Actinobacteria* (21%), *Planctomycetes* (6.25%), *Acidobacteria* (5.7%), *Bacteroidetes* (4%), *Firmicutes* (2.3%), *Verrucomicrobia* (1.5%) and *Cyanobacteria* (1%). The abundance levels of 5 phyla remained below 1%, namely, *Chloroflexi* (0.87%), *Gemmatimonadetes* (0.83%), *Nitrospirae* (0.28%), *Deinococcus*-*Thermus* (0.27%) and *Spirochaetes* (0.13%). Moreover, the abundance levels of 20 phyla remained below 0.1%, namely, *Armatimonadetes* (0.08%), *Chlorobi* (0.08%), *Chloridia* (0.06%), *Aquificae* (0.05%), *Thermotogae* (0.05%), *Ignavibacteriae* (0.05%), *Thermodesulfobacteria* (0.04%), *Kiritimatiellaeota* (0.04%), *Rhodothermaeota* (0.04%), *Deferribacteres* (0.03%), *Calditrichaeota* (0.02%), *Synergistetes* (0.02%), *Fusobacteria* (0.02%), *Tenericutes*(0.02%), *Candidatus Saccharibacteria* (0.01%), *Dictyoglomi* (0.01%), *Elusimicrobia*(0.001%), *Balneolaeota* (0.01%), *Atribacterota* (0.008%) and *Candidatus Omnitrophica* (0.01%). The least abundant phyla were *Chrysiogenetes* (0.009%), *Candidatus Bipolaricaulota* (0.009%), *Candidatus Cloacimonetes* (0.006%), *Fibrobacteres* (0.005%), *Coprothermobacterota* (0.004%), *Caldiserica* (0.003%) and *Candidatus Absconditabacteria* (0.0007%). The most abundant archaeal sequence was *Euryarchaeota* (0.24%), while the abundance levels of the other phyla did not exceed 0.1%, including *Thaumarchaeota* (0.05%), *Crenarchaeota* (0.03%), *Candidatus Thermoplasmatota* (0.008%), *Candidatus Lokiarchaeota* (0.002%), *Candidatus Korarchaeota* (0.0009%), *Candidatus Micrarchaeota* (0.0006%), and *Candidatus Nanohaloarchaeota* (0.0001%). A total of 222 reads (0.03%) were assigned to viral genomes, whereas unassigned and unclassified reads (2253067) accounted for 72% of the total reads on average (Fig.1B and 1C).

For fungi, 61,221 reads were assigned at the phylum level. The most abundant fungal sequence was *Ascomycota* (57%), followed by *Basidiomycota* (22%), *Mucoromycota* (8%), *Chytridiomycota* (6.9%), *Microsporidia* (1.2%) and *Zoopagomycota* (1.1%). At the genus level, *Aspergillus* (4.5%), *Rhizophagus* (2.7%), *Spizellomyces* (2.6%), *Batrachochytrium* (2%), *Lobosporangium* (2%), *Synchytrium* (1.8%), *Phycomyces* (1.5%), *Fusarium* (1.3%), *Exophiala* (1.3%) and *Rhizopus* (1.3%) were identified (Fig. 1E).

These genera represent the top 10 dominant fungal populations identified. Unassigned and unclassified reads (3048362) accounted for 98% of the total reads on average.

### Pantanal

At the phylum level, 748,195 and 8,303 reads were assigned to the *Bacteria* and *Archaea* domains, respectively. The most abundant bacterial sequence was *Proteobacteria*, accounting for 45% of the total sequences, followed by *Actinobacteria* (35%), *Planctomycetes* (3.7%), *Acidobacteria* (3.5%), *Bacteroidetes* (2.3%), *Firmicutes* (2.2%), *Chloroflexi* (1%) and *Gemmatimonadetes* (1%). The abundance levels of 5 phyla remained below 1%, namely, *Cyanobacteria* (0.9%), *Verrucomicrobia* (0.65%), *Nitrospirae* (0.5%), *Deinococcus*-*Thermus* (0.3%) and *Spirochaetes* (0.1%). Moreover, the abundance levels of 16 phyla remained below 0.1%, namely, *Chlorobi* (0.06%), *Armatimonadetes* (0.05%), *Aquificae* (0.05%), *Thermotogae* (0.05%), *Chlamydiae* (0.04%), *Thermodesulfobacteria* (0.04%), *Rhodothermaeota* (0.03%), *Ignavibacteriae* (0.03%), *Deferribacteres* (0.03%), *Kiritimatiellaeota* (0.02%), *Synergistetes* (0.02%), *Calditrichaeota* (0.02%), *Tenericutes* (0.02%), *Fusobacteria* (0.02%), *Candidatus Saccharibacteria* (0.01%), and *Balneolaeota* (0.01%). The least abundant phyla were *Elusimicrobia* (0.001%), *Atribacterota* (0.009%), *Dictyoglomi* (0.009%), *Candidatus Omnitrophica* (0.009%), *Candidatus Bipolaricaulota* (0.006%), *Chrysiogenetes* (0.006%), *Candidatus Cloacimonetes* (0.004%), *Caldiserica* (0.004%), *Fibrobacteres* (0.003%), *Coprothermobacterota* (0.003%), and *Candidatus Absconditabacteria* (0.001%). The most abundant archaeal sequences were *Thaumarchaeota* (0.74%) and *Euryarchaeota* (0.3%), whereas the abundance levels of the other phyla did not exceed 0.1%, including *Crenarchaeota* (0.03%), *Candidatus Thermoplasmatota* (0.01%), *Candidatus Korarchaeota* (0.01%), *Candidatus Lokiarchaeota* (0.001%), *Candidatus Micrarchaeota* (0.0006%), and *Candidatus Nanohaloarchaeota* (0.0004%). A total of 178 reads (0.02%) were assigned to viral genomes, whereas unassigned and unclassified reads (1523581) accounted for 67% of the total reads on average (Fig. 1B and 1C).

For fungi, 52,896 reads were assigned at the phylum level. The most abundant fungal sequence was *Ascomycota* (60%), followed by *Basidiomycota* (21%), *Chytridiomycota* (6.6%), *Mucoromycota* (6.4%), *Microsporidia* (1.2%) and *Zoopagomycota* (1.1%). At the genus level, *Aspergillus* (5%), *Fusarium* (3.4%), *Spizellomyces* (2.4%), *Batrachochytrium* (2%), *Talaromyces* (2%), *Lobosporangium* (1.9%), *Synchytrium* (1.8%), *Rhizophagus* (1.5%), *Phycomyces* (1.3%), and *Trichoderma* (1.3%) were identified. These genera represent the top 10 dominant fungal populations identified. Unassigned and unclassified reads (2227361) accounted for 97.5% of the total reads on average (Fig. 1E).

### General microbial community composition across biomes and microbiomes

To better understand the relationships between the different Brazilian biomes and the microbiomes (soil, rhizosphere and roots) within each biome, we created a nonmetric multidimensional scaling graph with all samples categorized by microbiome and biome (Fig. 2A). We employed the Bray-Curtis dissimilarity coefficient to measure the compositional dissimilarity between the genomes. We observed a greater correlation between samples from the same microbiome than between samples from the same biome. The root samples were grouped, as were the soil and rhizosphere samples. We observed a closer relationship between the soil and rhizosphere samples, with greater differentiation from the root samples (Fig. 2A). Notably, the Venn diagram yielded the same observation (Fig. 2B). Here, we observed that more operational taxonomic units (OTUs) were shared between the soil and rhizosphere samples, between the root samples, and between the soil or root samples and the rhizosphere samples. The dominant phyla in the root samples were cyanobacteria, which is true across all biomes (Fig. 3C). Given that the heatmap shown in Fig. 3C is organized by sample similarity, we observed a mixture of soil and rhizosphere samples dominated by *Proteobacteria*. Compared with the other biomes, the Caatinga exhibited a greater presence of *Firmicutes*. We generated rarefaction curves for our samples in two ways to assess the a diversity of the various biomes or microbiomes (Fig. 2D). As expected, the roots were less diverse than the soil and rhizosphere were (right). Biomewise, the Caatinga, PNB and Atlantic Forest exhibited very similar a diversity values at the lower end of the graph. Moreover, the Amazon and Pantanal exhibited similar but slightly greater diversity values, followed by Pampa. Notably, the Cerrado demonstrated the highest a diversity (Fig. 2D-left).

**Fig. 2.**
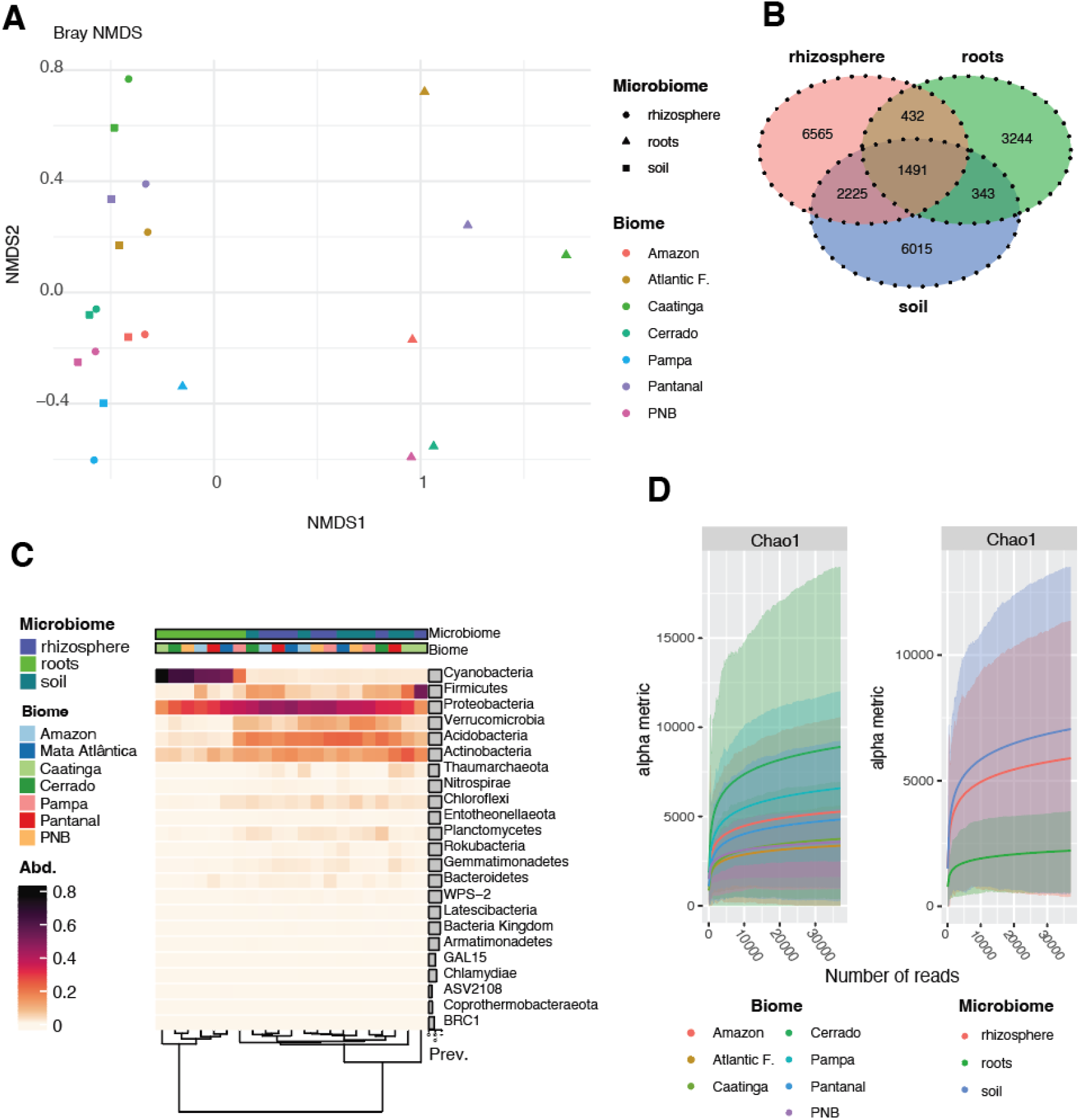
General profiling of the biome and microbiomes analyzed. **A** Non-metric multidimensional scaling (NMDS-Bray-Curtis) separated by biome and microbiome. Graph was made using the phyloseq package using the “plot_ordination” function. Best solution is as follows: Run 20 stress 0.09588036, Procrustes: rmse 4.94524e-06, max resid 1.675529e-05, Similar to previous best, Best solution repeated 9 times **B** Venn diagram of the OTUs shared between the microbiomes. Venn diagram was calculated and plotted in R using the VennDiagram package. **C** Heat-map of the Phylum of all samples separated by biome and microbiome. Heatmaps were made in R using the microViz package. **D** a-diversity using Chao index calculated with the function ggrarecurve from the MicrobiotaProcess R package.

**Figure 3.**
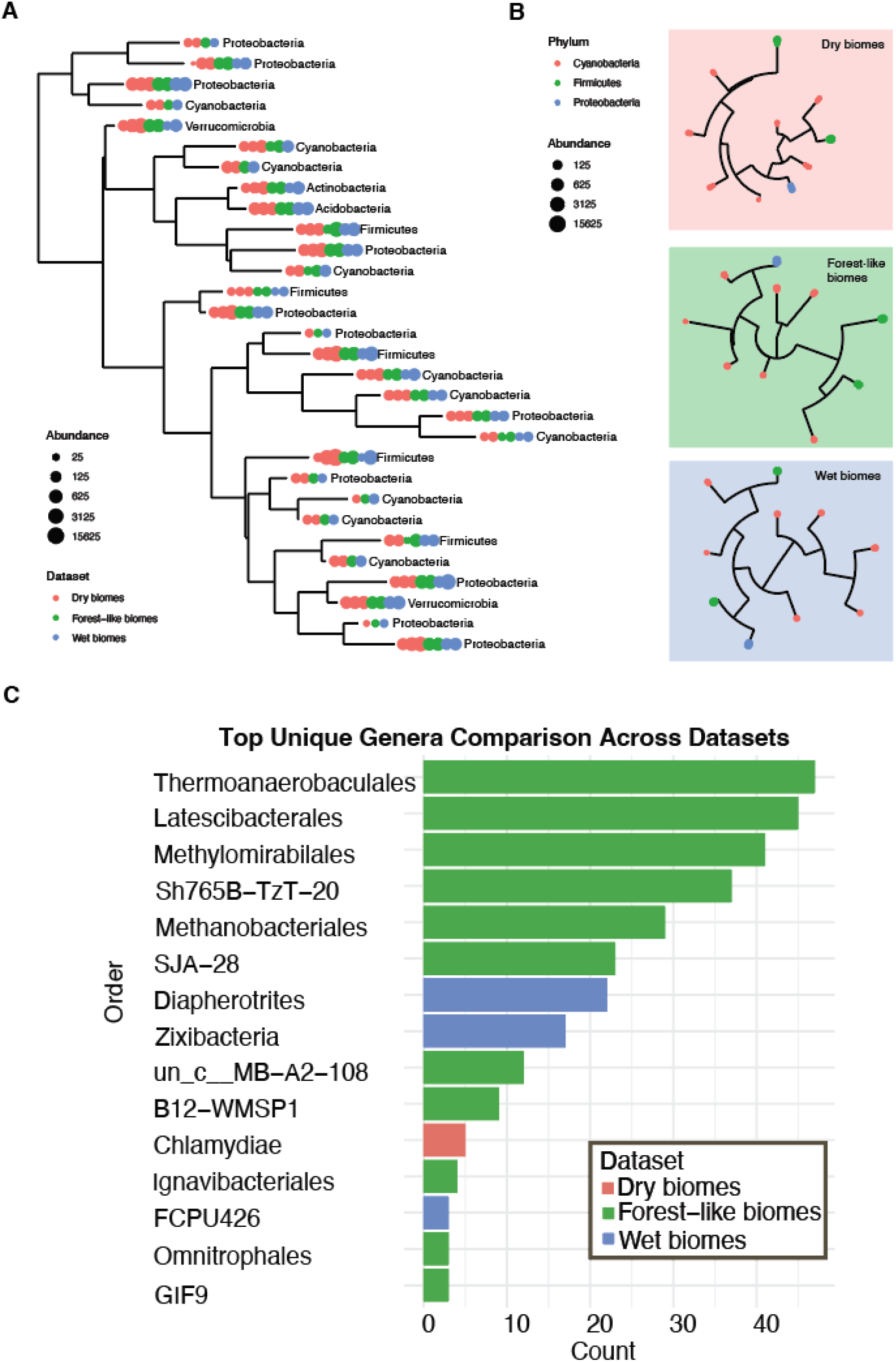
Metrics of dry, wet and forest-like biomes. **A** Taxonomic tree (Phylum) of all samples separated by Dry biomes, Forest-like biomes and Wet biomes. Plotted with Phyloseq function plot_tree **B** Phylum taxonomic trees of the biome categories shown separately. Plotted with Phyloseq function plot_tree **C** Unique top Orders of the different biome categories.

### The distribution of bacterial and archaeal phyla among Brazilian biomes exhibits significant diversity (differences between forested and dry biomes)

In our examination of the general diversity of microorganisms, the four most abundant phyla in the soil, root and rhizosphere samples from all the Brazilian biomes were *Proteobacteria, Actinobacteria, Bacteroidetes* and *Firmicutes* (Fig. 1B). These phyla include representatives that are easily isolated from cultures and widely studied. *Proteobacteria* was the most dominant bacterial phylum across almost all biomes, except the Caatinga. Surprisingly, *Actinobacteria* was especially prevalent in the semiarid Brazilian biome (Fig 1B and 1C). This notable shift in microbial dominance differs from the typical predominance of *Proteobacteria* in moist forest soils.

Furthermore, representatives of the *Acidobacteria* and *Planctomycetes* phyla were identified in the read annotations. *Planctomycetes* occur more frequently in the Cerrado and Pampa (Fig. 1C and 1D). *Acidobacteria* was more widespread in forested biomes such as the Amazon, Cerrado and Atlantic Forest biomes than in the Caatinga and Pantanal biomes (Fig. 1B). These findings highlight a distinct microbial community structure in the semiarid Caatinga environment compared with that in the other Brazilian ecosystems.

To better reveal the differences between the various Brazilian biomes, we separated them into forest-like biomes, wet biomes and dry biomes according to their general characteristics. Notably, we classified the Amazon and Atlantic Forest biomes as forest-like biomes. Moreover, we classified the Pampa and Pantanal biomes as wet biomes, as both are subject to flooding, while the Cerrado and Caatinga biomes were classified as dry biomes (Fig. 2A).

We plotted the top 10 unique orders in each category (Fig. 2B). Interestingly, in the wet biomes, we identified *Diapherotrites* and *Zixibacteria*, which are known to thrive in anaerobic environments such as wetlands, such as those in the Pantanal. The orders unique to the wet biomes are involved in nutrient cycling, organic matter degradation, and bioremediation of wet environments. The orders unique to the forest-like biomes included *Thermoanaerobaculales, Latescibacterales* and *Methylomirabilales*, which are generally associated with anaerobic environments that contain abundant organic matter, such as the Amazon and Atlantic Forest soil environments, which are rich in forest litter. We also assessed the relative abundance under each condition among the shared orders and between the three environments (Fig. 2D). The distribution of shared orders was similar among the three biomes. However, the order *Myxococcales* was absent in the dry biomes, as was the order *Oligoflexales* (Fig.2D).

### Comparison of the relative abundance levels of prokaryotic communities in the soil, roots and rhizosphere

Via the use of 16S amplicon-sequencing data, we distinguished the distribution patterns of phyla by separating the samples into soil, root and rhizosphere categories at all locations within the various Brazilian biomes (Fig. 3 and Supplemental Fig. 1). To visualize these trends, we generated bar plots depicting the richness of microbial phyla across the various biomes. This approach provides a more nuanced understanding of the difference in the composition of microbial communities among various ecological niches.

Notably, in the roots, cyanobacteria dominated, whereas the soil and rhizosphere were dominated by *Firmicutes* and *Proteobacteria*. An intriguing observation was obtained for outliers within these patterns. In the rhizosphere section, the Caatinga biome exhibited higher diversity than the other biomes and was characterized by a greater abundance of *Firmicutes* than *Proteobacteria*. Conversely, in the root section, the Pampa biome was identified as an outlier, featuring a lower abundance of cyanobacteria than that in the other biomes. In the soil section, the Cerrado–PNB sample from Cerrado–CV vegetation was noteworthy. Here, a distinct difference was observed, with a greater abundance of *Verrucomicrobia* than *Firmicutes*. This finding highlights unique microbial dynamics in the Cerrado–PNB, increasing our understanding of microbial diversity in Brazilian biomes.

### Comparison of the relative abundance levels of fungi in the soil, roots and rhizosphere

Via the use of sequence data from the 18S V4 and ITS-1 regions, we distinguished the distribution patterns of fungal and protist phyla by stratifying the samples from the different Brazilian biomes into soil, root and rhizosphere categories (Fig. 4A). This dual approach allowed us to compare the taxonomic diversity revealed by the conserved region (V4) with the specific variability captured by the ITS-1 spacer, highlighting how these complementary markers reflect the structure of microbial communities in distinct compartments of the soil-plant system. Comparative analysis of the most abundant fungal phyla in the Brazilian biomes revealed that *Ascomycota* was dominant in all environments, reaching a peak in the Amazon (0.077%), followed by Caatinga (0.068%), Cerrado–CV (0.043%), Pantanal (0.035%), Cerrado–PNB (0.057%), Atlantic forest (0.014%) and Pampa (0.015%), revealing a clear correlation between relatively high humidity and fungal abundance levels (Fig. 4B). *Basidiomycota*, the second most abundant phylum, reached a peak in the Amazon (0.061%), with decreasing values in the Cerrado–PNB (0.007%) and Pantanal biomes (0.005%), and minimum proportions in the other biomes (0.004–0.008%). The abundance levels of less representative phyla (*Mucoromycota, Zoopagomycota* and *Chytridiomycota*) generally did not exceed 0.01%, except *Mucoromycota* in the Amazon (0.008%) and Cerrado–PNB biomes (0.005%), suggesting that these groups occupy more specialized ecological niches (Fig. 4C). Interesting patterns emerged when comparing the humid biomes (Amazon, Atlantic Forest, and Pantanal) with the drier biomes (Caatinga, Cerrado, and Pampa). Notably, the former maintained more diverse and abundant fungal communities, probably due to the greater availability of organic matter and stable humidity conditions. Seasonal or arid biomes contained less diverse communities, but *Ascomycota* maintained its dominance, which suggests a remarkable adaptive capacity of this phylum. Moreover, ITS sequencing analysis revealed that the microbial compositions of the Brazilian biomes indicated unique genera, except those of the Caatinga and Cerrado biomes, which did not encompass unique genera. In the Amazon biome, the genera *Didymella, Xylogone*, order *Saccharomycetales*, Class *Chytridiomycetes* and *Filobasidium* were identified. The Cerrado-PNB biome exhibited *Lachnum*, order *Conioscyphales*, family *Leotiaceae, Thanatephorus* and *Olpidium*. The Atlantic Forest biome contained *Arachnotheca, Paecilomyces, Myriodontium, Lepiota* and *Cylindrocladium*, whereas the Pampa biome was characterized by *Auricularia, Vanrija*, family *Clavariaceae, Cryptococcus* and *Clavulinopsis*. In the Pantanal biome, *Acaulium, Pseudolophiostoma, Periconia, Arxiella* and *Angustimassarina* were identified as unique genera (Fig. 4D). These results highlight how abiotic (climate and humidity) and biotic (substrate availability) factors shape the structure of fungal communities at the biogeographic scale, with *Ascomycota* emerging as the most ubiquitous and resilient group in all the ecosystems analysed.

**Fig. 4.**
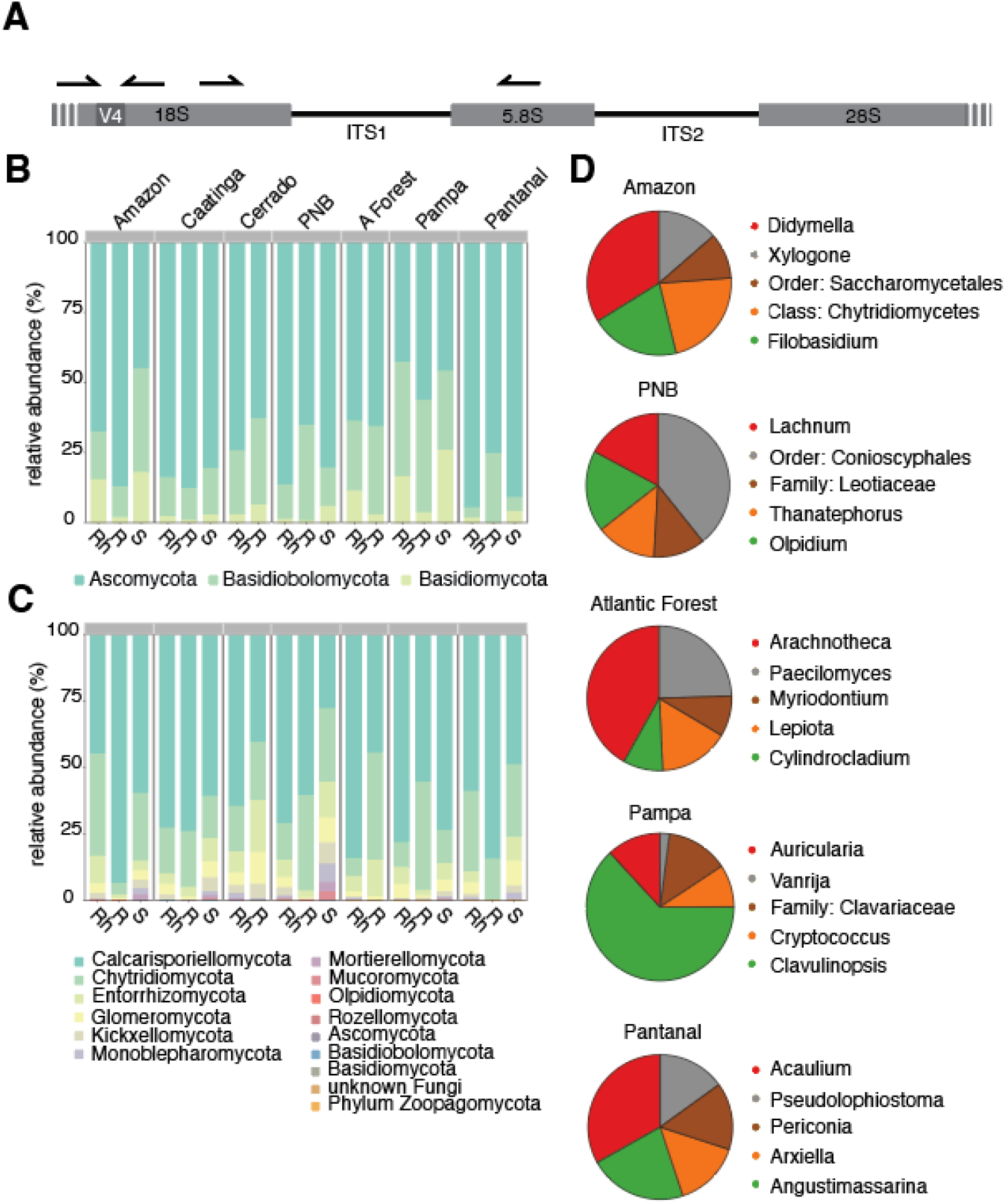
ITS classifications of the Eukaryotic microbes from the Brazilian biomes. **A** Location of the primers used to amplify the 18S and the ITS regions that were sequenced. **B** Top three most abundant phylum from the Biomes. Rh refers to rhizosphere, R refers to roots and S refers to soil samples. **C** After the removal of the top 3 most abundant phyla, remaining phyla were plotted. **D** Top 5 unique genus from ITS sequencing of the Brazilian biomes. Caatinga and Cerrado did not have any Genus that was unique to the biome.

## Discussion

The diversity of the soil, root and rhizosphere microbial communities in representative areas of all Brazilian biomes was assessed. *Proteobacteria* dominated the bacterial communities in nearly all the biomes. This phylum includes bacteria involved in carbon, sulfur and phosphate cycling and symbiotic nitrogen fixation (SNF), such as *Bradyhizobium, Burkholderia, Paraburkholderia* and *Cupriavidus [44-50]*. This finding can be linked to the high levels of soil organic matter (SOM) and moisture in soils, which favour fast-growing heterotrophs, particularly those involved in nitrogen cycling [51, 52]. The second most abundant phylum was *Actinobacteria*, such as *Streptomyces*, which are involved in various processes, such as phosphate solubilization, nitrogen cycling, and organic matter catabolism [53, 54].

In the Amazon, *Acidobacteria* and *Planctomycetes* were well represented, both of which are known to encompass representatives that thrive in acidic and oligotrophic environments [55, 56]. With respect to archaea, *Nitrososphaera* and *Candidatus Nitrosocosmicus* (*Thaumarchaeota*) dominated the community, and they are often associated with ammonia oxidation in soils, suggesting a key role in nitrogen and nutrient cycling in Amazonian soil processes [57, 58].

The Atlantic Forest exhibited greater seasonal variations in moisture and temperature than the Amazon did [59, 60]. Therefore, while *Proteobacteria* still dominated *Actinobacteria*, its proportion was greater. Moreover, *Thaumarchaeota* and *Euryarchaeota* remained the most abundant archaea with a similar pattern to that in the Amazon, further highlighting the importance of nitrogen cycling across forested biomes [57, 58, 61].

The distribution of microbial phyla between CV and PNB in the Cerrado exhibited a similar distribution pattern but with interesting differences. *Proteobacteria* continued to dominate in both Cerrado areas, with a notable presence of *Actinobacteria*. However, differences emerged because *Actinobacteria* was more abundant in PNB than in CV. This difference may be explained by the fact that the Cerrado is characterized by a pronounced dry season, which suggests that microbial communities can adapt to fluctuating moisture levels [62]. Furthermore, CV contained twice as many *Planctomycetes* as did PNB. These phyla thrive in oligotrophic environments, which could suggest nutrient limitations in Cerrado soils [56]. *Acidobacteria* and *Firmicutes* exhibited lower yet significant variations between the two regions, with CV containing more *Acidobacteria* and *Firmicutes* than PNB does, potentially indicating differences in the soil pH, as *Acidobacteria* are sensitive to acidic conditions [55, 63]. The archaeal community was less diverse in the Cerrado than in the Amazon, with *Euryarchaeota* as the most abundant group.

Pampa is similar to temperate grasslands, where nitrogen fixation and decomposition are critical processes [8, 64, 65]. Thus, *Proteobacteria* and *Actinobacteria* continued to dominate, but *Bacteroidetes* were more abundant than in the Amazon and Cerrado. This phylum includes bacteria involved in the degradation of complex polysaccharides, which could be linked to the greater input of grassland plant material [66, 67]. However, archaea were dominated by *Euryarchaeota*, reinforcing their widespread ecological role in methanogenesis and nitrogen/phosphate/carbon cycling.

### The Pantanal exhibited a notable shift in *Actinobacteria*

High water availability and seasonal flooding might explain the relatively high levels of *Actinobacteria* and *Proteobacteria*, which play significant roles in soil organic matter decomposition and nutrient cycling. Furthermore, *Planctomycetes* and *Acidobacteria* demonstrated relatively low abundance levels, possibly reflecting the relative dynamic water and nutrient influx conditions. In addition, the most abundant archaeal groups were *Thaumarchaeota* and *Euryarchaeota*. The slightly greater proportion of *Thaumarchaeota* may indicate a significant contribution of ammonia-oxidizing archaea to nitrogen cycling in wetland soil.

In the Caatinga, the relative abundance of *Proteobacteria* was lower than that in the Amazon and Atlantic Forest, reflecting the lower availability of nutrients and moisture, which limits the fast-growing heterotrophic members of this phylum. In addition, *Actinobacteria* accounted for nearly half of the total bacterial sequences, which was significantly more abundant than in the other biomes. This suggests high specialization for drought-tolerant microorganisms due to the arid conditions in the Caatinga because the ability of *Actinobacteria* to tolerate desiccation and their role in organic matter decomposition are likely key to maintaining ecosystem function in semiarid and arid regions [68-70]. Moreover, archaeal diversity was characterized by high proportions of *Thaumarchaeota* and *Euryarchaeota*, suggesting that despite the low-nutrient and low-moisture conditions, archaea still play an essential role in nitrogen and carbon cycling.

Fungi are essential soil components that mediate indispensable processes in nutrient cycling, such as decomposers, saprophytes, symbionts and parasites [71, 72]. *Rhizophagus* species are AMF, a group of roots obligate biotrophs with symbiotic relationships with plant roots [73]. They are considered natural biofertilizers that help plants absorb essential nutrients such as phosphorus and nitrogen from soil. The distribution of fungi across Brazilian biomes revealed that *Ascomycota* was the most abundant phylum in all the biomes. This phylum is known for its diverse ecological roles and ability to adapt to distinct environments, likely contributing to its prevalence [74-76]. *Aspergillus* was consistently the most abundant genus in all the biomes, indicating its adaptability and widespread distribution in various ecological niches. *Trichoderma* and *Fusarium*, known for their roles in plant symbiosis and pathogenic interactions, occurred prominently in the Pantanal and Atlantic Forest. *Trichoderma* exhibits high potential for use in preventing diseases, promoting plant growth, enhancing nutrient utilization efficiency, increasing plant resistance and mitigating agrochemical pollution [77, 78]. *Fusarium* encompasses several phytopathogenic species that cause great economic losses worldwide. In general, they produce a wide variety of mycotoxins, and the consumption of products contaminated with mycotoxins can cause acute or chronic effects in animals and humans and can result in immunosuppressive or carcinogenic effects [79].

### *Basidiomycota* followed *Ascomycota* in terms of fungal read abundance

These fungi fulfil significant roles in the degradation of lignin-rich plant litter and are therefore associated with nutrient cycling and decomposition, which are vital in soil ecosystems [80, 81]. Other phyla, i.e., *Mucoromycota, Chytridiomycota, Microsporidia* and *Zoopagomycota*, exhibited relatively low but uniform proportions across the various biomes. *Mucoromycota* was more abundant in Pampa and Pantanal biomes, and *Chytridiomycota* was more abundant in the Pampa and Caatinga biomes than in the other biomes. These differences, even if slightly greater, may indicate that these fungi may be better suited to grassland ecosystems with more temperate and variable conditions. *Microsporidia* maintained a stable, low-level presence in all biomes. Despite its relatively low abundance, this phylum is an important parasitic group that can impact microbial population dynamics, potentially contributing to the overall complexity of soil microbial communities [82, 83]. Ultimately, despite being the minor fungal phylum in terms of abundance, *Zoopagomycota* maintained a stable presence across all biomes. This phylum includes fungi that prey on small soil organisms such as other fungi, amoebae and nematodes and may play a niche-specific role in microenvironments where competition for resources is high, thereby helping to modulate microbial dynamics in nutrient-limited soils and extreme conditions such as drought (semiarid) or flooding (floodplain) [84, 85].

Fungi is involved in plant symbiosis and stress tolerance fulfils crucial roles in increasing plant survival and productivity. Biomes with nutrient-poor or extreme conditions, such as the Caatinga, Pantanal and Cerrado (CV and PNB) biomes, exhibited relatively high levels of fungi involved in plant symbiosis and stress tolerance. *Aspergillus* was abundant across all the biomes.

Differences in microbial distribution are influenced by soil characteristics and preferences for specific host plants or organisms. Furthermore, the quality and scope of the samples collected directly influence the representativeness of the microbial diversity results. Environmental factors must also be considered, as they can affect the observed diversity, and the analysis must account for the specific conditions in the ecosystems studied. Each biome supports microbial communities that are ecologically specialized to local conditions. Studying the dynamic forest x microbiome in an integrated manner is essential for understanding microbial interactions, ecosystem functions, and responses to global changes across diverse habitats [86]. Forests favour nutrient cycling in acidic soils, whereas drier biomes promote decomposers and microorganisms capable of surviving in harsh nutrient-poor environments. In general, the distributions of minor phyla in all biomes provide insights into local ecological differences. For example, low-abundance phyla exhibit slight shifts, suggesting that environmental factors, such as temperature, light exposure, soil pH, moisture availability, and organic matter content, and anthropogenic factors, such as land use and agricultural practices, may shape their presence and niche opportunities. Additionally, spatial and ecological factors, such as elevation, landscape structure, soil composition and vegetation cover, likely lead to the creation of distinct microhabitats, thus influencing the microbial composition at each location.

## Conclusion

The distribution of microorganisms in the soil, rhizosphere and roots across Brazilian biomes reflects the complex interactions among environmental factors such as moisture, nutrient availability, and soil properties. The dominance of certain phyla in specific biomes suggests a specialized ecological role for these microorganisms, from nitrogen fixation in the Amazon to soil organic matter decomposition in the Caatinga. Moreover, the consistent presence of *Thaumarchaeota* across biomes, particularly in the Amazon and Pantanal, suggests that ammonia oxidation by archaea plays a role in nitrogen cycling, especially in soils with varying nitrogen availability. Future research could focus on exploring the functional roles of these microbial communities in more detail, providing insights into how these ecosystems might respond to environmental changes such as climate change and land-use shifts. Future studies should investigate the functional resilience of biome-specific microbial communities under simulated stress and functional metagenomics, while also exploring multi-trophic interactions between drought-adapted microbiota and native vegetation to unlock their biotechnological potential in sustainable agriculture and medicine.

## Figures

Figures were made in Adobe illustrator. References to the package used to calculate the figures in R are described in each figure’s legend.

## Data availability

All data generated or analyzed during this study are included in this published article and its supplementary information files. Correspondence and requests for materials should be addressed to Elibio Rech.

## Acknowledgments

We would like to thank Claudomiro de Almeida Cortês from the Association of native seed collectors of Cerrado da Chapada dos Veadeiros for his support in defining collection sites. We also thank Luís Henrique Mota de Freitas Neves, Maria Carolina Alves de Camargos and Alexandre Bonesso Sampaio from Chapada dos Veadeiros National Park for the legal authorizations and support in defining collection sites.

We deeply thank the Chico Mendes Institute for Biodiversity Conservation - ICMBio (authorizations 85243 and 85247) for the support and legal authorizations granted for the collection of soil samples.

We also acknowledge the National Institute of Science and Technology in Synthetic Biology, National Institute of Science and Technology in Engineering Biological Systems and the Ministry of Agriculture and Livestock. Funding from the National Council for Scientific and Technological Development (465603/2014-9; 400145/2023-5; 308815/2023-8), Research Support Foundation of the Federal District (0193.001.262/2017), and the Coordination for the Improvement of Higher Education Personnel.

## Author contributions

L.M.A.T., R.N.L. and E.R. designed the study; C.A.X.A., G.M.S.R., D.M.C.B., D.S., M.S.B.M., J.P.P.T., L.B.S.V., F.A.F., R.L. and J.L.S.A conducted field sampling; L.M.A.T. and R.N.L. performed laboratory experiments; L.M.A.T., R.N.L., P.V.P., M.A.O., D.B., P.M., M.F. performed data analysis; E.R. secured funding.

All authors completed, edited and approved the final version.

