## Supplemental figure 1 for "Bridging Gaps in Soil Ecology: Metagenomic Insights into Microbial Diversity and Functionality Across Brazil’s Biomes"

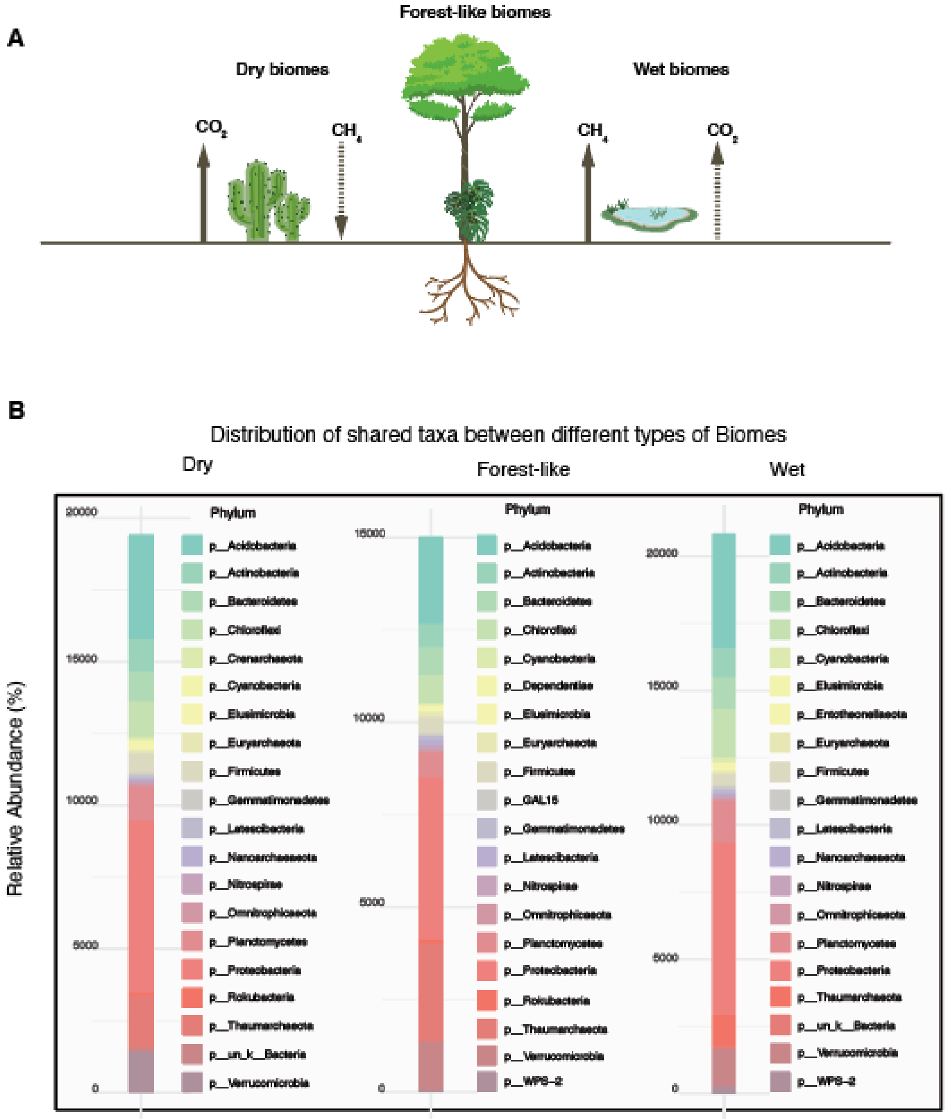


Supplemental figure 1. Differences between dry, wet and forest-like biomes. A. Graphical representation of the three different biome categories. B. Distribution of the shared phylum taxa on their relative abundances for each biome category.
